# Exonuclease assisted mapping of protein-RNA interactions (ePRINT)

**DOI:** 10.1101/2023.05.16.540978

**Authors:** Sophie Hawkins, Alexandre Mondaini, Seema C. Namboori, Asif Javed, Akshay Bhinge

**Affiliations:** College of Medicine and Health, University of Exeter, EX1 2LU, UK; Living Systems Institute, University of Exeter, Ex4 4QD, UK; School of Biomedical Sciences, LKS Faculty of Medicine, The University of Hong Kong, Hong Kong SAR, China

## Abstract

RNA processing is a fundamental mode of gene regulation that is perturbed in a variety of diseases including cancer and neurodegenerative disorders. RNA-binding proteins (RBPs) regulate key aspects of RNA processing including alternative splicing, mRNA degradation and localization by physically binding RNA molecules. Current methods to map these interactions, such as CLIP, rely on purifying single proteins at a time. We have developed a new method (ePRINT) to map RBP-RNA interaction networks on a global scale without purifying individual RBPs. ePRINT allows precise mapping of the 5’ end of the RBP binding site, and can uncover direct and indirect targets of an RBP of interest. Importantly, ePRINT can also uncover RBPs that are differentially activated between cell fate transitions, for instance, as neural progenitors differentiate into neurons. Given its versatility, ePRINT has vast application potential as an investigative tool for RNA regulation in development, health and disease.

## Main

Physical interactions between RNA binding proteins (RBPs) and RNAs regulate key aspects of cellular homeostasis including RNA biogenesis, splicing, transport, localization, and decay [1]. Disruption of these regulatory interactions leads to cellular dysfunction and disease [2]. Several RBPs have been implicated in human diseases including cancers and neurodegenerative disorders [3,4]. Current gold-standard methods to map RBP targets involve biochemical purification of the RBP-bound RNA followed by sequencing of the RNA cargo [5,6]. This necessitates availability of high-quality antibodies to perform the RBP-RNA purification which excludes a majority of RBPs expressed in mammalian cells. RBPs regulate thousands of genes including those encoding RBPs within intricate interaction networks. Mapping these networks on a global scale requires immunopurification of hundreds of RBPs, a massively expensive and laborious task, especially if the goal is to compare changes in RBP-RNA networks across cell states (for example disease vs healthy). To circumvent these issues, we developed exonuclease-assisted mapping of protein-RNA interactions (ePRINT), a new method that allows mapping RBP-RNA interactions across the transcriptome without the need to purify specific RBPs. Our method exploits the recent observation that organic extraction of UV cross-linked cell lysates causes RBP-RNA complexes to migrate to the interphase [7]. We enrich these RBP-RNA complexes and sequence the RNA to identify the footprint of the bound protein (Fig.1A). Using bioinformatic analyses, we then uncover the identity of the RBP at each locus and map changes in RBP activity between experimental conditions.

**Figure 1.**
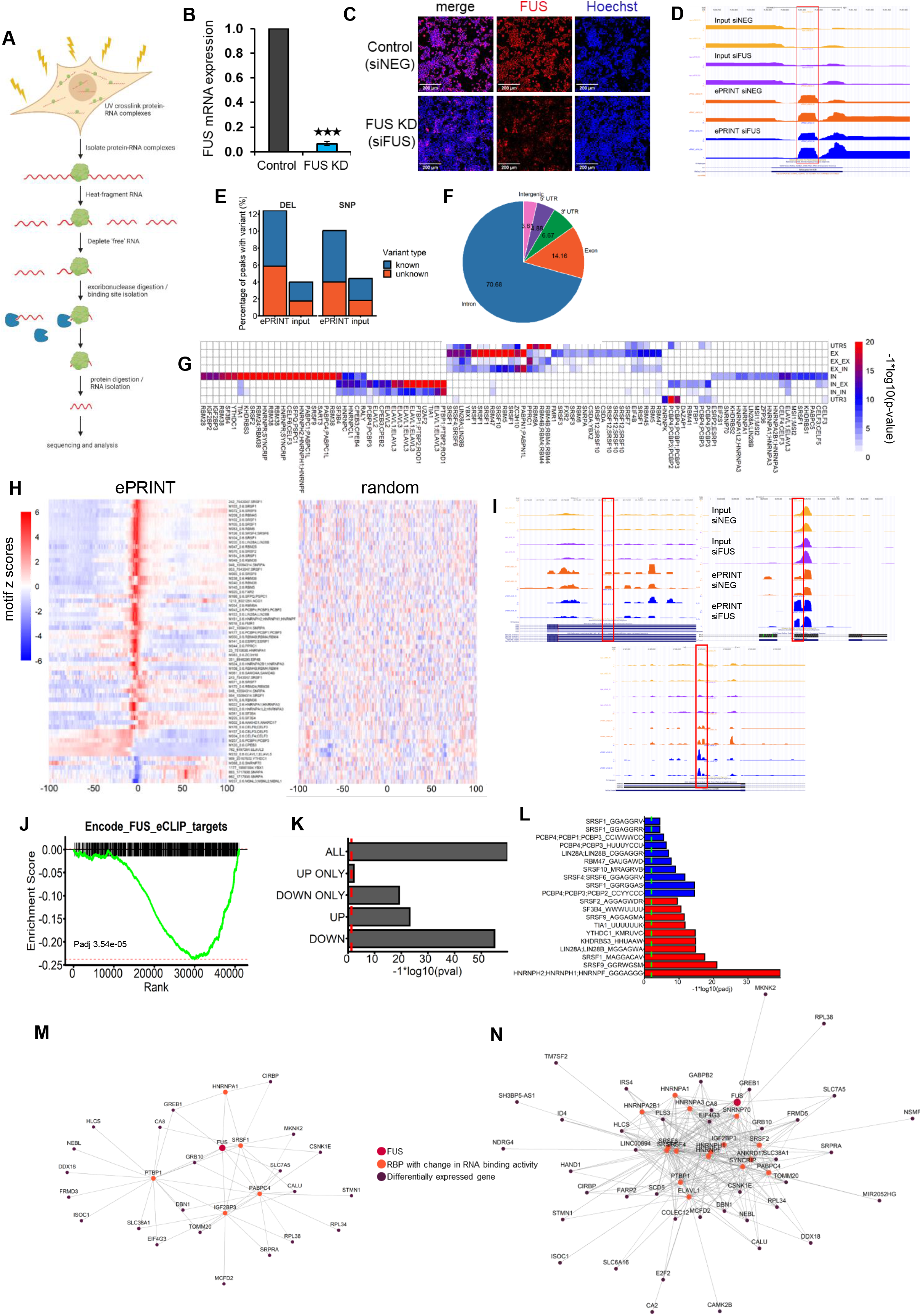
ePRINT identifies bonafide RBP-RNA interactions. **<u>A)</u>** Schematic of the ePRINT protocol. Briefly, cells are cross-linked using UV irradiation then lysed. Protein-RNA complexes are isolated, then RBP-RNA binding sites are isolated by heat fragmentation followed by 5’-3’ exonuclease digestion. Finally, the protein is digested and the RNA encoding the protein footprint is sequenced. **<u>B)</u>** RT-qPCR indicating siRNA-mediated depletion of FUS mRNA in HEK293T cells after 72 hours. N = 4. **<u>C)</u>** Representative images indicating siRNA-mediated depletion of FUS protein in HEK293T cells after 72 hours. **<u>D)</u>** UCSC genome browser snapshot showing an example RBP peak in exon 4 of the XIST gene that is enriched in ePRINT vs input, and unchanged between experimental conditions. **<u>E)</u>** Mutational analysis showing percentage of peaks with deletions (DEL) or single nucleotide polymorphisms (SNPs). Known variant indicates that this mutation is found within the 1000 genomes database [33]. **<u>F)</u>** Distribution of all ePRINT peaks identified in HEK293T cells across the following gene features: Introns, Exons, 5’ UTRs, 3’UTRs and intergenic regions. Numbers indicate percentages of ePRINT peaks mapped to a given feature. **<u>G)</u>** Enrichment of RBP motifs in peaks mapping to different intragenic gene features. UTR5, EX, IN, and UTR3 indicate peaks where both the start and end sites map within the same 5’UTR or exon or intron or 3’UTR respectively. EX_EX indicates peaks where the start and end sites map to different exons. EX_IN indicates peaks where the start and end sites map to exons and introns respectively. IN_EX indicates peaks where the start and end sites map to introns and exons respectively. IN_IN indicates peaks where the start and end sites map to different introns. **<u>H)</u>** RBP motifs are enriched at peak start sites indicated by 0 on the x-axis. Peak start sites were extended by 100bp (+/-). Scores for individual motifs were estimated at each bp along each peak using the position weight matrices. Per bp scores were averaged across all peaks and converted into z-scores: higher z-scores (red) indicate higher probability of locating the motif(s). Left panel indicates peaks identified in ePRINT samples. Right panel indicates randomly generated peaks. **<u>I)</u>** UCSC genome browser snapshots showing example RBP peaks that are altered in the siFUS condition compared to the siNEG control. <u>DOWN peak (upper panel)</u>: Region: 3’ UTR of the ITM2C gene. Log2FC - 8.64, padj 5.02e-07. <u>UP peak (middle panel):</u> Region: Exon 6 of the RPSA gene. Log2FC 2.47, padj 4.65e-09. <u>Adjacent UP/DOWN peak (lower panel):</u> Region: Region: 3’ UTR of the SMARCC1 gene. UP peak: Log2FC = 1.41. Padj = 5.90e-13. DOWN peak: Log2FC = -0.86. Padj = 4.35e-04. **<u>J)</u>** Peak set enrichment analysis of FUS binding sites identified using eCLIP in HepG2 (GSE177328) and K562 (GSE176874) cell lines. X-axis indicates ePRINT peaks ranked from most significantly upregulated (left side) to most significantly downregulated (right side). Padj indicates the Benjamini-Hochberg adjusted p-value. **<u>K)</u>** Hypergeometric test to determine enrichment of FUS targets** in genes that display enhanced (UP) or reduced (DOWN) ePRINT peaks upon FUS knockdown. Genes with peaks in both directions are excluded in the UP ONLY and DOWN ONLY comparisons. **<u>L)</u>** Top 10 RBP motifs identified as enriched in peaks that are enhanced (red) or reduced (blue) after FUS knockdown. An enhanced peak indicates that the associated RBP has more binding events, or is more active, after FUS knockdown. A reduced peak indicates that the associated RBP has fewer binding events. **<u>M,N)</u>** Network analysis to identify direct and indirect effects of FUS knockdown. FUS CLIP datasets GSE177328 and GSE176874 were used to identify direct targets of FUS. eCLIP datasets (M) or ePRINT peaks (N) were then used to identify targets of the RBPs showing a change in activity from figures 1L and S.7B. eCLIP captures 23/53 of the genes that are differentially expressed upon FUS knockdown (DEGs). ePRINT captures 40/53 DEGs. **FUS targets are a consolidation of 4 FUS CLIP datasets with the following geo accessions: GSE177328, GSE176874, GSE118347 and GSE50178. *** indicates pval <0.001 by Student’s t-test. Error bars indicate SEM

We performed organic extraction on UV crosslinked HEK293T cells and quantified the amount of RNA retained in the aqueous phase. As expected, increasing UV strength led to a decrease in the levels of aqueous phase RNA with an UV dose of 400 mJ/cm2 recovering ∼74% of the total RNA from the interphase (Fig.S1A). Having confirmed that covalently bound protein-RNA complexes can be effectively purified, we sought to identify the RNA footprint occupied by the bound protein. We heat fragmented the RNA before organic extraction (Methods) to a median size of 100-200 nucleotides (Fig.S1B) and purified the protein-RNA complexes from the interphase. As expected, crosslinked lysates allowed efficient recovery of fragmented RNA while non-crosslinked lysates showed a significant loss of material (Fig.S1B). However, we observed recovery of low amounts of fragmented RNA from the non-crosslinked samples (Fig.S1B). To get rid of the observed background, we treated the recovered RNA with T4 polynucleotide kinase to introduce a 5’ phosphate and then digested the repaired RNA with the exonuclease XRN1. We hypothesized that XRN1 activity will cause complete digestion of the unbound RNA while covalently bound protein will physically occlude XRN1 progression, enabling precise mapping of the 5’ end of the protein footprint. XRN1 digestion further reduced RNA amounts from both the non-crosslinked and cross-linked samples with significantly more RNA being recovered from the cross-linked sample (Fig.S1B). However, we were unable to completely remove RNA from the non-crosslinked sample. This could be due to a subset of the RNAs being resistant to XRN1 digestion, possibly due to the 5’ end of the RNAs being protected by a 5’ cap or incomplete repair leading to retention of a 5’ hydroxyl group. Subsequently, we digested the RNA-bound protein with Proteinase K to release the RNA and prepared Illumina compatible libraries for sequencing using the NEBNext small RNA kit. Our library preparation protocol requires adapter ligation to the 3’ and 5’ ends of the RNA. Hence, any RNA fragments with protected 5’ ends were not expected to be included in the final library. In parallel, we purified 1/10th of each cross-linked sample without performing any enrichment to be used as a RNA fragment size-matched input.

We applied ePRINT to detect global RBP-RNA interaction networks regulated by the RBP FUS. FUS is localised to the nucleus and is involved in regulating RNA splicing as well as microRNA biogenesis [8]. It is also found to be dysregulated in multiple cancers as well as neurodegenerative disorders including Amyotrophic Lateral Sclerosis [4,9]. Given its importance in cellular homeostasis, multiple studies have explored the downstream targets of FUS using CLIP making it an attractive choice to validate ePRINT. We knocked down FUS using siRNAs in HEK293T cells (Fig.1B, Fig.1C), and performed ePRINT across two biological replicates. We detected 204736 peaks that were identified within genes in at least two out of the four samples (Methods). Analysis of individual binding events across genes showed the importance of using an un-enriched size matched input to filter out background read densities in ePRINT experiments (Fig.S2). We only retained peaks that were significantly enriched relative to the size-matched input generating a total of 41567 high-confidence RBP-RNA binding events (example peak, Fig.1D).

Analysis of CLIP reads has revealed an enrichment of deletions or substitutions at the UV cross linked protein-RNA interaction sites due to errors in reverse transcription over the cross-linked nucleotides. [10,11]. Analysis of the reads overlapping the ePRINT peaks uncovered a significant enrichment of deletions and substitutions in reads derived from the ePRINT samples compared to input (Fig.1E). Interestingly, not all substitutions were enriched equally. We noticed that A>G or T>C substitutions were more commonly observed than G>A and C>T (Fig.S3). This could indicate a preference of A and T nucleotides to be crosslinked to proteins compared to G or C. Further, the identified ePRINT sites overlapped significantly more with at least one ENCODE eCLIP peaks compared to randomly selected sites in the transcriptome (29.84% in ePRINT vs 10% in random simulations, p-value < 2.2 × 10e-6) indicating that ePRINT was identifying true binding events. Distribution of the ePRINT peaks across gene features revealed an abundance of sites in introns (Fig.1F, Fig.S4A), in accordance with the role of RBPs in alternative splicing and mRNA stability. Next, we performed motif enrichment analysis to uncover the identities of the RBPs that potentially bind specific gene features (Fig.1G). Strikingly, motifs enriched in introns were assigned to RBPs known to function in the nucleus including ELAVL3, SFPQ and several HNRNP proteins, indicating that the identified proteins bind pre-spliced mRNAs in the nucleus. These motifs were significantly depleted in other gene features such as exons and UTRs (Fig.S4B). On the other hand, motifs enriched in exons or UTRs belonged to RBPs known to be involved in translational regulation or mRNA localization including LIN28 and FMR1 (Fig.1G). As expected, such motifs were depleted in the introns indicating these RBPs function in the cytosol (Fig.S4B). Mapping motif locations relative to peak start coordinates revealed that most motifs were enriched close to the start site (Fig.1H). This indicates that XRN1 digestion continues until it is impeded by the protein-RNA crosslinking site allowing precise mapping of the crosslinked location.

Next, we utilised ePRINT data to identify genome-wide FUS binding sites. We expected that knockdown of FUS would lead to a decrease in ePRINT peak amplitude at FUS binding sites. We identified differentially bound protein-RNA sites using a statistical model that normalises the change in ePRINT peak amplitude to the change in host gene expression after FUS knockdown (Methods). Differential analysis identified 2753 peaks as increasing in amplitude (enhanced peaks) and 2403 as decreasing in amplitude (reduced peaks) in response to FUS knockdown (example peaks in Fig.1I). We sorted the peaks from most enhanced to reduced and compared the sorted list of peaks with previously published eCLIP-identified FUS binding sites using gene set enrichment analysis (GSEA). GSEA identified a significant enrichment of the FUS eCLIP peaks in the reduced ePRINT peakset (Fig.1J, Fig.S5A), confirming our expectation. We further validated the overlap between ePRINT and eCLIP datasets at the gene level, allowing us to incorporate additional FUS CLIP datasets in our analysis. We first mapped all FUS CLIP peaks within intragenic regions to the corresponding gene to define direct FUS targets. Next, we applied the same criteria to differential ePRINT peaks, and calculated the overlap between the ePRINT and CLIP gene sets by a hypergeometric test. The overlap was calculated separately for the set of genes that showed enhanced or reduced ePRINT peaks. We noticed that a subset of genes displayed both enhanced and reduced peaks within the gene body. To avoid confounding our results, we also performed the overlap analysis after removing such genes. Gene targets of ePRINT peaks that were reduced after FUS knockdown showed the most significant overlap with FUS eCLIP gene targets (Fig.1K, Fig.S5B), with the enrichments at par when comparing FUS targets across cell lines (Fig.S6). Strikingly, the overlap at the gene level returned highly significant p-values compared to the overlap at the peak level. This suggests that FUS has multiple closely aligned binding sites per gene and each CLIP dataset might be capturing only a subset of these sites. Overall, our analysis indicates that ePRINT can identify FUS binding events across the whole transcriptome with high confidence. We searched the set of reduced peaks for motifs previously associated with FUS binding. We observed an enrichment of GU-rich motifs that were shown to be enriched in previously published FUS CLIP datasets [12–14](Fig.S7A). However, we also identified enrichment of GU-rich motifs in upregulated peaks indicating that GU enriched motifs are not specific to FUS binding (Fig.S7A). Previous studies on FUS motif analysis have noted a lack of consistency in assigning a sequence motif to FUS [15]. It is possible that FUS uses a range of motifs for target recognition depending on the cell type.

Since ePRINT, in-principle, can capture all RBP-related binding events, we used the data to evaluate changes to the entire RBP-RNA interaction network in response to FUS knockdown. We first identified other RBPs that may be affected by FUS knockdown by performing motif analysis on the differentially bound ePRINT peaks (Methods). The reduced set of peaks was found to be enriched for the Serine and Arginine Splicing Factor RBPs (Fig.1L, Fig.S7B) in line with the expected role of FUS in regulating alternative splicing. Motif analysis also retrieved the RBP FMR1 (Fig.S7B), which is known to act synergistically with FUS [16]. Accordingly, we observed an enrichment of a FMR1 binding sequence in the set of downregulated peaks. Our analysis indicates that knockdown of FUS impairs FMR1 binding to cognate RNAs. On the other hand, upregulated peaks were enriched for the HNRNP proteins HNRNPH, HNRNPF and HNRNPC, and TIA1 (Fig.1L, Fig.S7B). FUS belongs to the HNRNP family, sometimes known as hnRNP P2 [17]. Our analysis indicates that in the absence of FUS, these RBPs might expand their target repertoire to regulate FUS targets, potentially as a compensatory mechanism.

FUS depletion in HEK293T cells resulted in the differential expression of 53 genes (FUS KD DEGs) (supplementary table 1). Analysis of published CLIP data suggested that FUS directly interacts with only 5 of these, indicating that the remaining 47 genes could be regulated by RBPs that are downstream from FUS. From the list of RBPs identified from our motif analysis that showed differential binding upon FUS knockdown, we identified the RBPs that are direct FUS targets (FUS-RBPs) using FUS CLIP data. Next, we used the ENCODE CLIP datasets to determine which of the 53 FUS KD DEGs are direct targets of our FUS-RBPs. CLIP datasets were available only for 5 out of the 16 FUS-RBPs. Using this approach, we were able to link 23/53 of the FUS KD DEGs to at least one of the FUS-RBPs (Fig.1M). We repeated this analysis using our ePRINT data to link the FUS KD DEGs to FUS-RBPs. As we were no longer limited by the availability of CLIP datasets, we were able to analyse all candidate RBPs that are directly regulated by FUS. This resulted in linking 40/53 FUS KD DEGs to at least one FUS RBP (Fig.1N). Thus, using ePRINT, we were able to generate a more comprehensive network of the indirect effects of FUS depletion as compared to using available CLIP datasets.

Next, we evaluated the efficacy of ePRINT in determining RBP-RNA network changes across cell state transitions. The differentiation of self-renewing stem cells into neural progenitors and post-mitotic neurons is regulated by an intricate network of RBPs [18–20]. We deployed ePRINT to identify key RBPs regulating the transition of motor neuron progenitors (MNPs) into motor neurons (MNs). OLIG2+ MNPs were derived from human iPSCs and further differentiated into OLIG2-/ISL1+ MNs by pharmacologically inhibiting NOTCH signalling [21] (methods). Immunostaining for OLIG2, ISL1 and NFM indicated that MNPs and MNs were generated at high efficiency using our protocol (Fig.2A). ePRINT and peak finding was performed as described above with minor modifications (methods). We did not observe any significant changes in the distribution of peaks across gene features between the MNP and MN ePRINT samples (Fig.2B). Differential peak analysis using DESeq2 identified 889 peaks as enriched in MNPs and 5123 peaks enriched in MNs. Example peak changes can be seen in Fig.2C. Motif analysis on the differentially expressed peaks identified RBP motifs specifically enriched in MNPs or MNs (Fig.2D, Fig.S8). We mapped motif locations with respect to the peak coordinates and observed an enrichment of the motif around the peak start site for several top candidates (Fig.S9, Fig.S10). We decided to focus our attention on the RBP motifs that displayed a sharp enrichment at the peak start site, and that were predicted to drive MNP self-renewal (Fig.2E). We hypothesised that overexpression of RBPs responsible for maintaining MNPs in a state of self-renewal would inhibit neuronal differentiation. We delivered constructs encoding the RBPs HNRNPF and SRSF9 to MNPs and then induced them to differentiate using NOTCH inhibition (Fig.S11). Strikingly, over-expression of both candidate RBPs resulted in a significant reduction in neurite outgrowth compared to our control condition (Fig.2F, Fig.S12). Although both genes show downregulation at the transcript levels from MNP to MN (Fig.2G, Fig.2H), neither rank highly in the list of 163 RBPs which are differentially expressed between cell states (Supplementary Table 2). This highlights the power of ePRINT in identifying RBPs in terms of their activity and not simply a change in expression between cell states, a feature that will be instrumental for understanding RNA regulation in development, health and disease.

**Figure 2.**
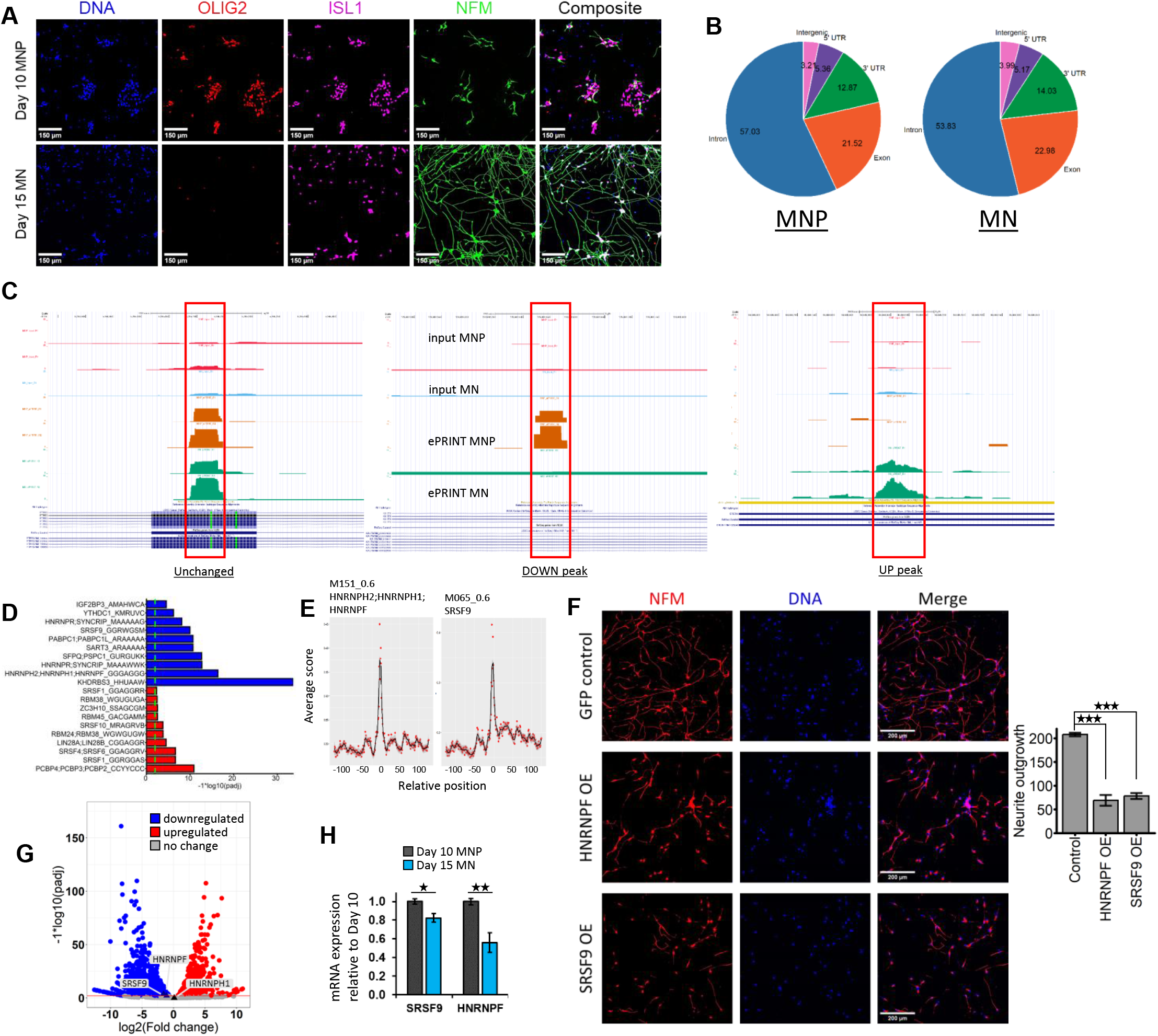
ePRINT uncovers RBPs regulating cell fate transition from motor neuron progenitors (MNPs) to post-mitotic motor neurons (MNs). **<u>A)</u>** Representative images of day 10 motor neuron progenitors (MNPs) and day 15 post-mitotic motor neurons (MNs). **<u>B)</u>** Distribution of ePRINT peaks across gene features in MNPs vs MNs. Numbers indicate percentages of ePRINT peaks mapped to a given feature. **<u>C)</u>** UCSC genome browser snapshots showing example RBP peaks. <u>Unchanged peak (left panel</u>): Region: Exon 5 of the PTPRS gene. Log2FC 4.86e-03, padj 1 (0.9999719). <u>DOWN peak (middle panel</u>): Peak that was identified as reduced in MNs compared to MNPs. Region: Intron 1 of the KALRN gene. Log2FC -8.76, padj 8.21e-05. <u>UP peak (right panel</u>): Peak that was identified as enhanced in MNs compared to MNPs. Region: Exon 19 of the SRCIN1 gene. Log2FC 12.45, padj 6.46e-07. **<u>D)</u>** Top 10 RBP motifs identified as enriched in peaks that are enhanced (red) or reduced (blue) in MNs compared to MNPs. **<u>E)</u>** Motifs M151_0.6 (HNRNPH2, HNRNPH1, HNRNPF) and M065_0.6 (SRSF9) display enrichment at the peak start site (indicated as 0 on the x-axis). Peak start sites were extended by 100 bp on either side for the motif analysis. **<u>F)</u>** Representative images of day 16 MNs after RBP overexpression (OE) (left panel) and quantification of total neurite outgrowth normalised by cell count (right panel). Control indicates eGFP expression. Cells were stained with NFM to define the soma and neurites. Nuclei were stained using Hoechst (blue). **<u>G)</u>** Volcano plot of differentially expressed genes from day 10 MNP to day 15 MN. SRSF9, HNRNPF and HNRNPH1 have been annotated (light grey labels; triangle points). HNRNPH2 was not expressed in either MNPs or MNs. **<u>H)</u>** RT-qPCR analysis of SRSF9 and HNRNPF mRNA expression levels between day 10 MNPs and day 15 MNs. N = 3. * indicates pval <0.05. ** indicates pval <0.01. *** indicates pval <0.001 by Student’s t-test. Error bars indicate SEM

To further ascertain the mechanism behind the observed phenotypes, we sought to identify the downstream targets of these RBPs. To accomplish this, we mapped the motifs of these RBPs to ePRINT peaks upregulated in MNPs vs MNs. The identified peaks for each RBPs were then mapped to protein coding genes that were deemed as downstream targets of the said RBPs. Based on this analysis, we identified 995 peaks mapping to 857 genes for SRSF9 and 695 peaks mapping to 623 genes for HNRNPF. Gene ontology enrichment analysis on these genesets uncovered pathways targeted by these RBPs (Fig.S13). HNRNPF targets were enriched for cell-matrix adhesion, which are highly active in differentiation, embryonic development and remodelling events [22]. SRSF9 targets were enriched for morphological and developmental processes, alongside histone methylation. This suggests that SRSF9 may directly regulate proteins involved in cell morphology, but also affect transcription of neuronal (or non-neuronal) genes by affecting chromatin accessibility.

ePRINT simultaneously estimates transcriptome wide RBP binding thereby capturing a comprehensive picture of the role of all RBPs under a given condition and time point. However, that comes with the limitation of reliance on motif-based analytics to tease apart the roles of individual RBPs. In this study, we used 174 motifs mapping to 142 RBPs from the transite database [23]. These numbers are expected to increase with advancements in motif detection and the availability of high-quality CLIP datasets. Our method improves on two recent papers that also aim to map transcriptome wide RBP binding events [24,25]. ePRINT does not rely solely on organic extraction to enrich RBP-RNA complexes as this alone is insufficient to deplete unbound RNA. The addition of the exonuclease digestion not only increases the signal-to-noise ratio but also allows precise mapping of RBP-binding sites without the requirement of mutational analysis, allowing high confidence assessment of RBP motifs. Further, ePRINT can be applied for post-mortem tissue analysis, as live cells are not a requirement. Our analysis approach enables global identification of RBPs with a change in activity between cell states, whereas previous methods have focused on RBP profiles on a small subset of lncRNAs in a given cell state. Finally, by mapping ePRINT peaks to gene features, our analysis is accurately able to predict localisation of RBPs (Fig.1G, Fig.S4B). This could be instrumental in uncovering mislocalisation events often observed in neurodegenerative diseases such as ALS.

In summary, ePRINT can be deployed to identify downstream targets of a single RBP or map changes in the RBP interactome on a global scale as cells transition from one state to another in a cost-effective manner. With the increasing evidence implicating RBPs in a variety of diseases including neurodegeneration and cancer, we expect ePRINT will be a powerful method to understand the mechanistic basis of how RBP networks are altered in diseases.

## Data availability

The raw reads and bedgraph files for each sample have been deposited to the Gene Expression Omnibus database (GSE230097).

## Declaration

None of the authors have any competing interests.

## Methods

### 293T maintenance

HEK293T cells were maintained in Dulbecco’s modified Eagle’s medium (DMEM; Merck) supplemented with 10% fetal bovine serum (Merck), 2 mM GlutaMAX (Gibco) and 10 mM Hepes (Gibco).

### FUS knockdown

HEK293T’s were seeded at a density of 200,000 cells per 6-well. The next day, cells were transfected with 5nM Silencer Select siRNA (Thermo Fisher; siNeg, siFUS) using calcium-phosphate (Takara). Media was changed at 24 hours and samples were collected 72 hours post-transfection.

### Immunofluorescence

Cells were fixed in 4% paraformaldehyde, permeabilized in ice-cold methanol and blocked in 10% serum (HEK293T) or 1% BSA (MN). Primary antibodies (FUS 1:100 Santa Cruz Biotech sc-47711; ISL1 1:500 Abcam ab109517; OLIG2 1:100 R&D Systems AF2418; NFM 1:1000 Merck MAB1621) were added in 1% BSA and incubated at 4degC overnight. The next day, wells were washed with PBS, then incubated with Alexa-Flour secondary antibodies (Molecular probes; 1:2000) and Hoechst 33542 (Molecular probes, 1:1000) at RT for 1 hour. Cells were imaged using the ImageXpress Pico (Molecular Devices), or DMi8 microscope (Leica). Image quantification was completed using Cellprofiler [26].

### RT-qPCR

Cells were lysed and RNA extracted using the Monarch Total RNA Miniprep Kit (NEB) or RNA cleanup kit (NEB) according to manufacturer instructions. cDNA was reverse transcribed using random hexamers and the High Capacity reverse transcription system from Applied Biosystems. Quantitative PCR was performed using the SYBR GREEN PCR Master Mix from Applied Biosystems and the target gene mRNA expression was normalized to the expression of 2-4 housekeeping genes (HPRT1, RPL13, GAPDH and ACTB). Relative mRNA fold changes were calculated by the ΔΔCt method. Primer sequences are included below:

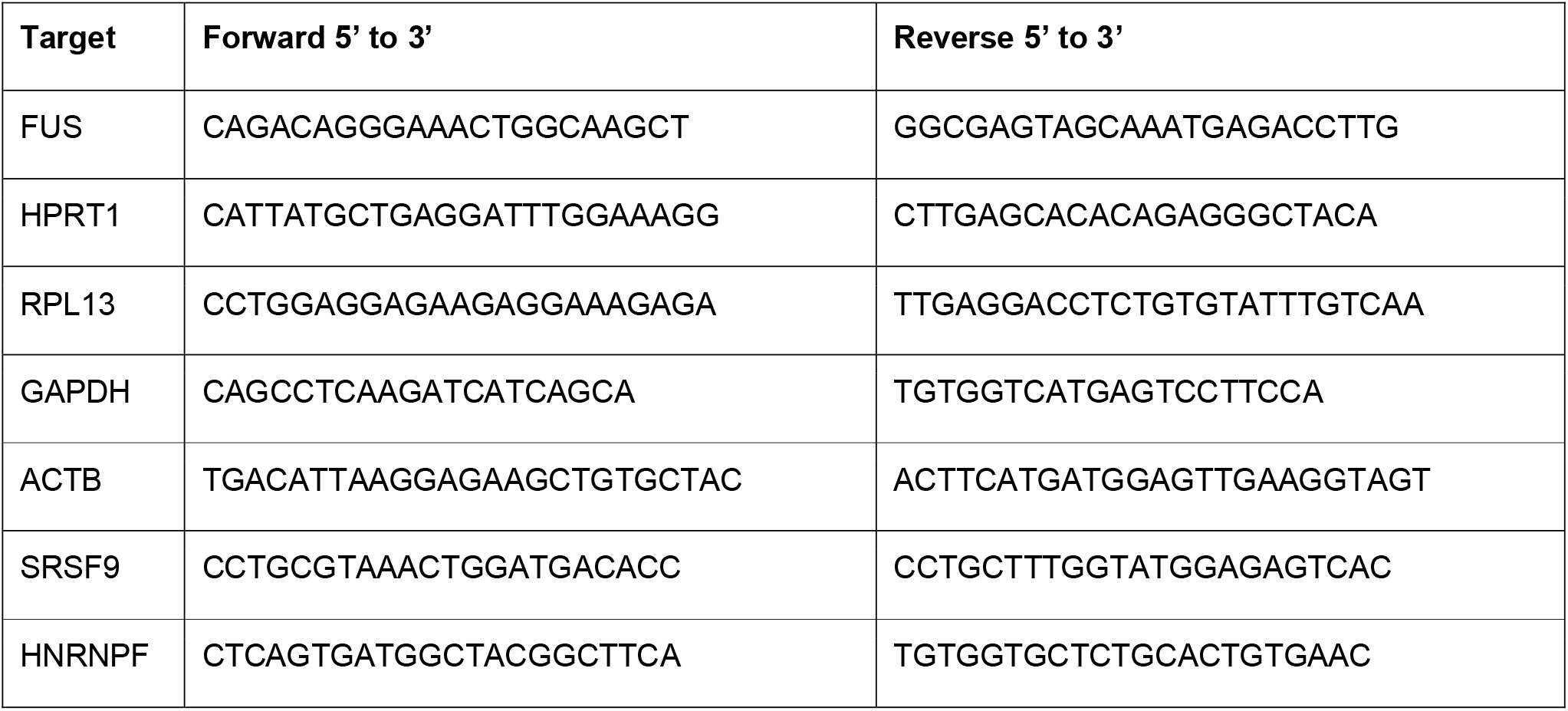

### ePRINT

Please contact Akshay Bhinge or Sophie Hawkins for the protocol.

### iPSC maintenance and MN differentiation

Healthy human iPSCs (GM23279A) were obtained from the Coriell Institute for Medical Research. iPSCs were maintained as colonies on human ES qualified Matrigel (Corning) in StemFlex (StemCell Technologies). Colonies were routinely passaged using EDTA and mycoplasma testing was conducted regularly to rule out contamination of cultures.

For differentiation to MN, iPSCs were plated as colonies onto Matrigel and treated with neuronal differentiation media (DMEM/F12:Neurobasal in a 1:1 ratio, HEPES 10mM, N2 supplement 1%, B27 supplement 1%, L-glutamine 1%, ascorbic acid 5uM) supplemented with SB431542 (40uM), CHIR9921 (3uM) and LDN8312 (0.2uM) from day 0 till day 4. Cells were caudalized by treatment with 0.1uM retinoic acid starting at day 2 and ventralized with 1uM purmorphamine starting at day 4 and continued till day 8. On day 8, motor neuron progenitors (MNPs) were re-plated onto poly-D-lysine/laminin coated wells. Differentiation was induced by treating the cells with N2B27 media supplemented with retinoic acid, purmorphamine and DAPT 10uM. DAPT treatment was stopped at day 13 and media was changed to N2B27 supplemented with BDNF 10μg/ml, GDNF 10μg/ml. Samples were collected at day 10 (MNP) and day 15 (MN) for ePRINT.

### RBP overexpression

HNRNPF, SRSF9 and eGFP mRNA sequences were inserted into the inducible pLV-TetO vector, replacing NGN2. Viral particles were produced using the pspax2 packaging vector and the pMD2.G envelope vector. The tet-encoding virus pLV_hEF1a_rtTA3 was generated in parallel. MNPs were transduced at day 9 and RBP expression was induced with 2 μg/mL doxycycline (dox) at day 10 (Fig.S11). Notch inhibition was conducted at day 11 using DAPT as described above and maintained until day 16 when cells were fixed for immunofluorescence. 24 hours prior to fixation, cells were selected with 500 ng/mL puromycin. Dox-containing media was replenished every 48 hours.

p3x-FLAG-hnRNPF was a gift from Mariano Garcia-Blanco, addgene #21926 [37]. SRSF9_2xRRM_pGEX was a gift from Christopher Burge, addgene #135120 [38]. pLV-TetO-hNGN2-Puro was a gift from Kristen Brennand addgene #79049 [39]. psPAX2 and pMD2.G were gifts from Didier Trono (Addgene plasmid #12260 and #12259). pLV_hEF1a_rtTA3 was a gift from Ron Weiss addgene #61472 [40].

