## Supplemental data for "Exonuclease assisted mapping of protein-RNA interactions (ePRINT)"

Figure S1

**A**

| UV mJ/cm2 | µg RNA in aqueous phase |
|---|---|
| 0 | 9 |
| 200 | 6 |
| 400 | 2.7 |
| **UV mJ/cm2** | **µg RNA in interphase** |
| 400 | 7.8 |

**B**

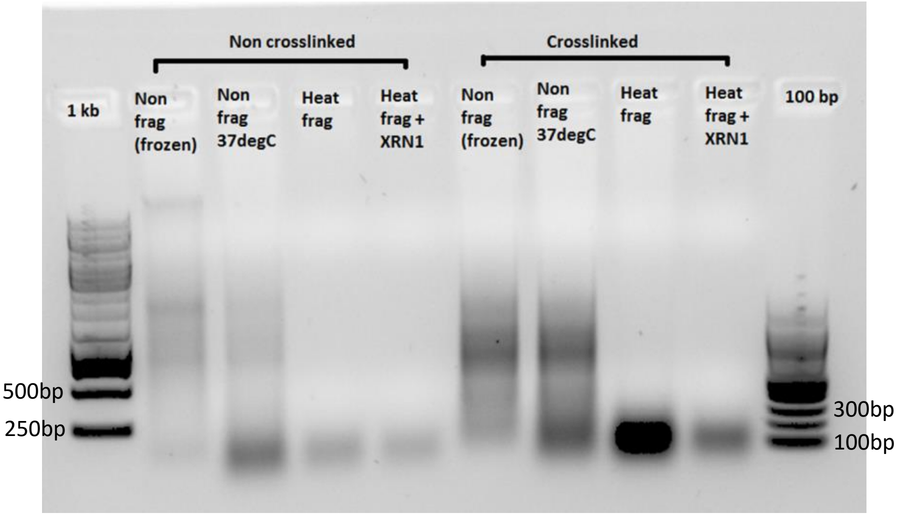


**Figure S1. ePRINT experimental optimizations**

**A).** RNA is depleted from the aqueous phase upon UV irradiation, and can be recovered from the interphase. In brief: HEK293T cells were irradiated at 0, 200 and 400 mJ/cm$^2$, then lysed and phase separated using QIAzol and chloroform. For the aqueous phase, free RNA was extracted using spin column purification. For the isolated interphase, protein was digested using proteinase K, then the RNA was isolated using spin column purification with gDNA removal. Both the aqueous and interphase extractions included DNase treatments.

**B)** RNA abundance isolated from interphase is increased upon UV irradiation (400 mJ/cm$^2$). Heat fragmentation for 30min, 94degC is optimal to reduce RNA size to ~100-200bp. XRN1 greatly reduces RNA abundance in crosslinked cells, indicating successful 5'-3' digestion.

### Figure S2

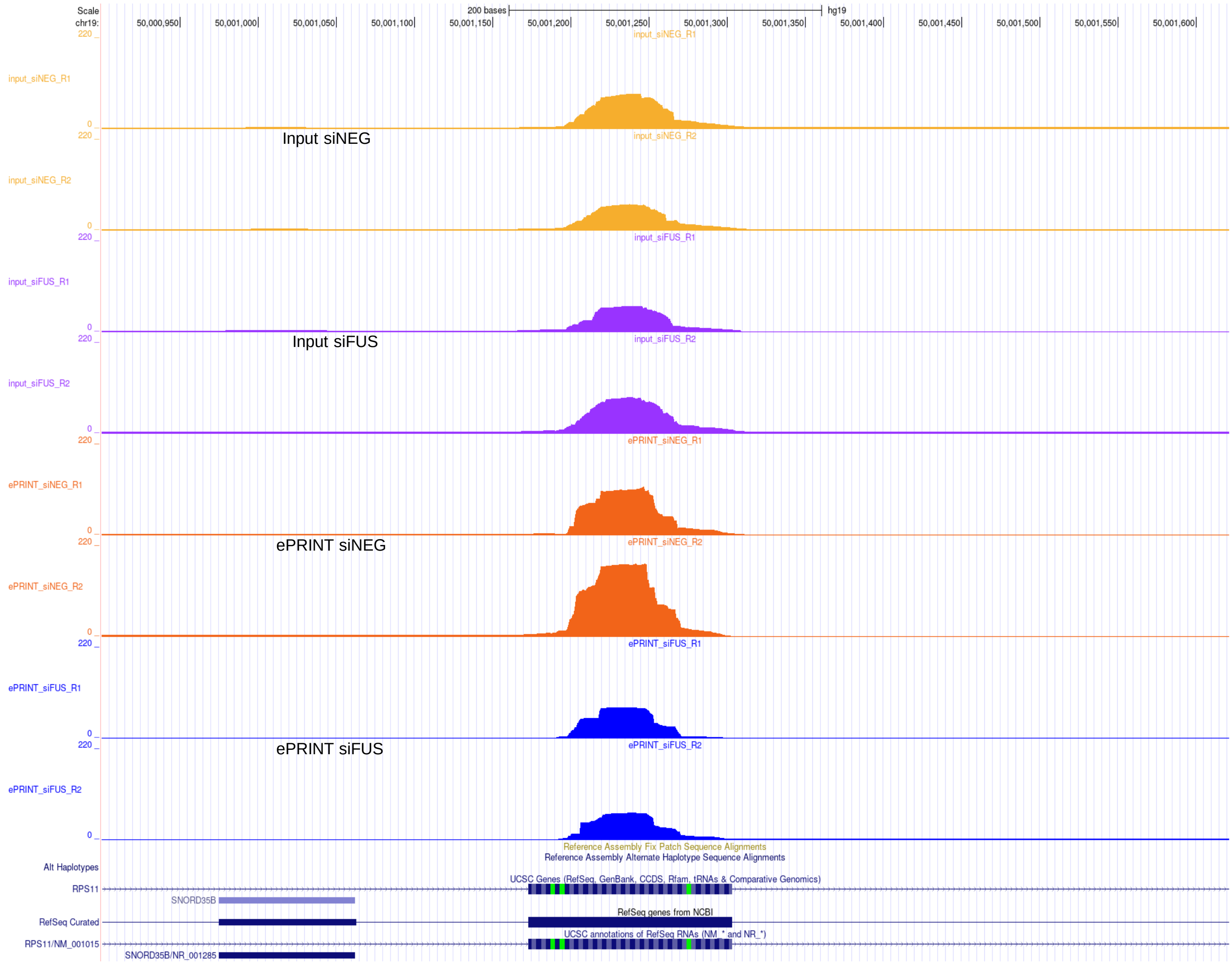


**Figure S2.** Representative UCSC browser screenshot showing the importance of input filtering to remove peaks that are not due to RBP binding events. Region: exon 4 of the RPS11 gene

### Figure S3

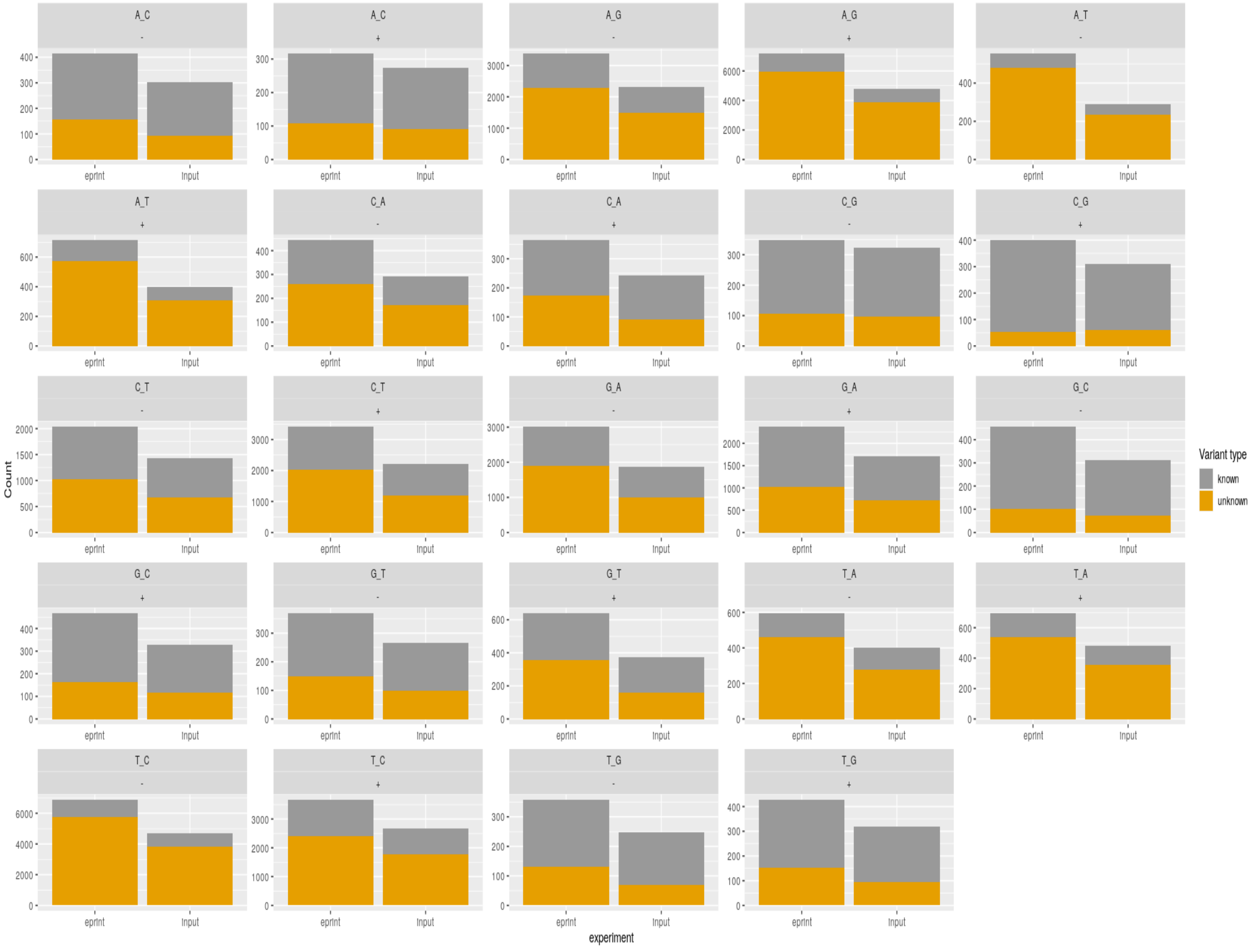


**Figure S3.** Mutational analysis identified base changes under ePRINT peaks compared with input. Related to Fig.1E

### Figure S4

A

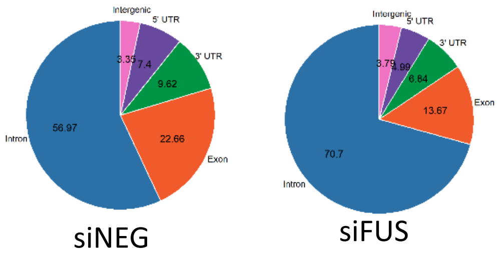


B

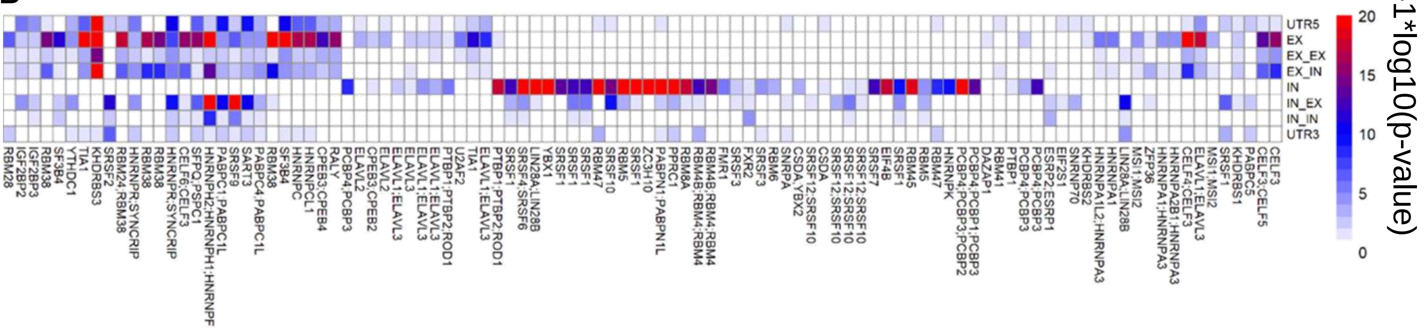


**Figure S4. ePRINT binding site distribution across gene features**

**A)** Distribution of ePRINT peaks mapped to gene features. Numbers indicate percentages of ePRINT peaks mapped to a given feature. Venn diagrams display data for siNEG and siFUS conditions. Associated with Fig.1F.

**B)** RBP motifs identified as depleted in specific gene features by hypergeometric test. Associated with Fig.1G.

### Figure S5

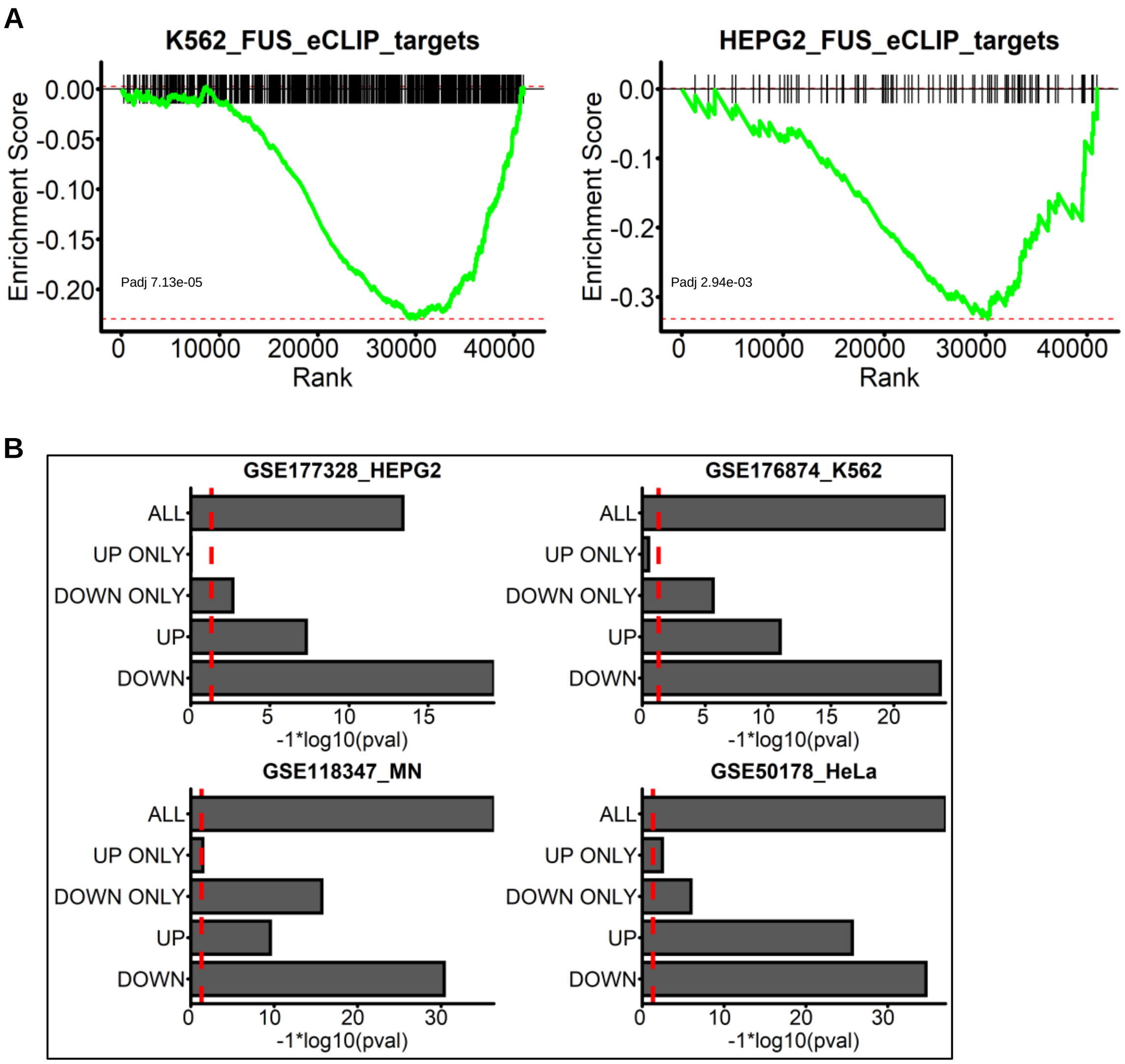


**Figure S5. Overlapping analysis of genes identified using ePRINT and eCLIP**

**A)** Individual peak set enrichment analysis of FUS binding sites identified by the encode eCLIP datasets GSE177328 (HepG2) and GSE176874 (K562). Associated with Fig.1J.

**B)** Overlap analysis to determine enrichment of FUS targets in genes that display enhanced (UP) or reduced (DOWN) ePRINT peaks upon FUS knockdown. Genes carrying peaks in both directions are excluded in the UP ONLY and DOWN ONLY comparisons. Associated with Fig.1K

### Figure S6

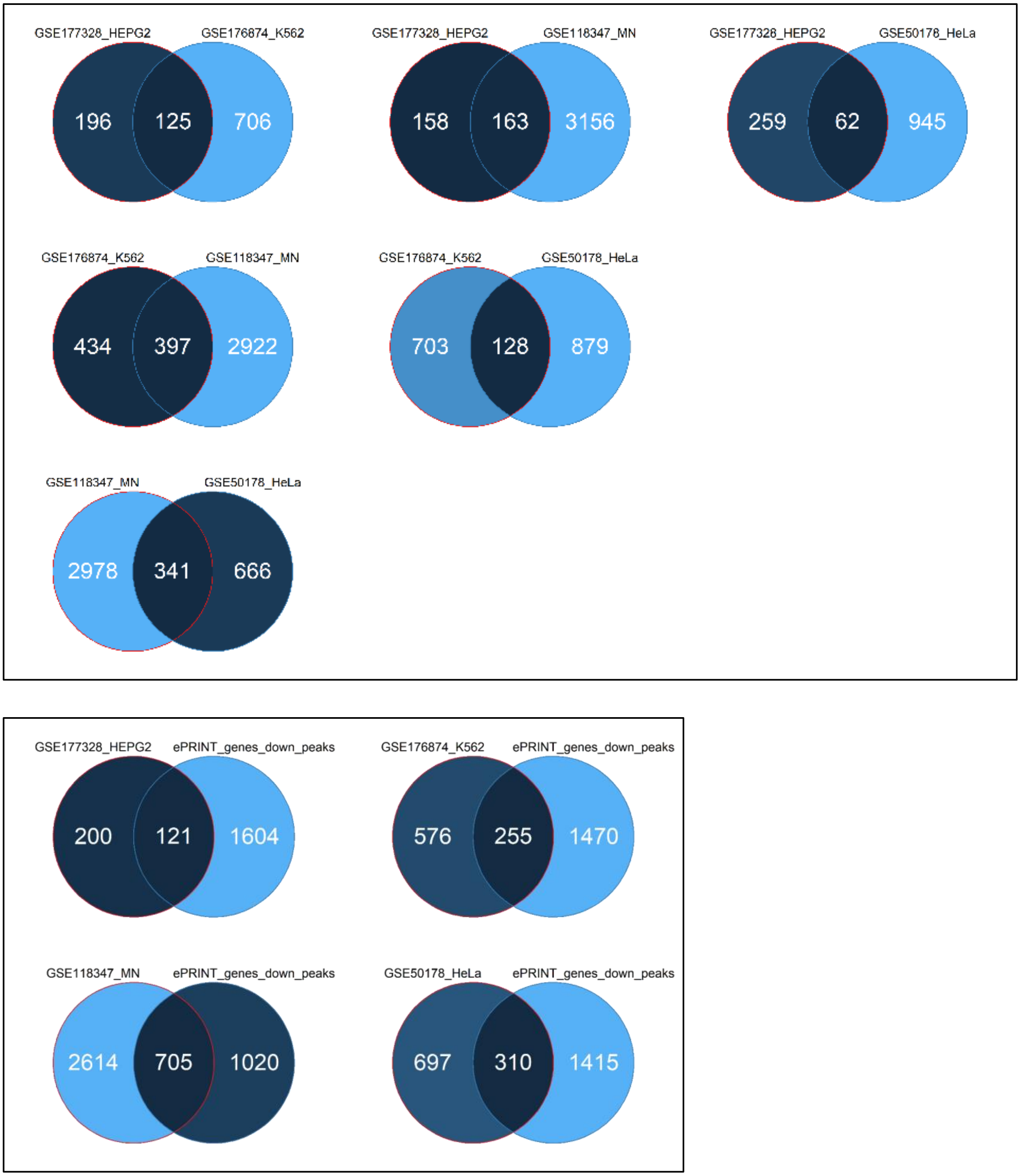


**Figure S6. Pairwise analysis of FUS target genes between CLIP datasets and ePRINT.**
**Top panel:** pairwise overlap of FUS gene targets identified by the four CLIP datasets used to generate figures 1K and S2E. **Bottom panel**: pairwise overlap of FUS CLIP datasets with genes identified by ePRINT carrying peaks that are redcued after FUS knockdown in HEK293T cells.

### Figure S7

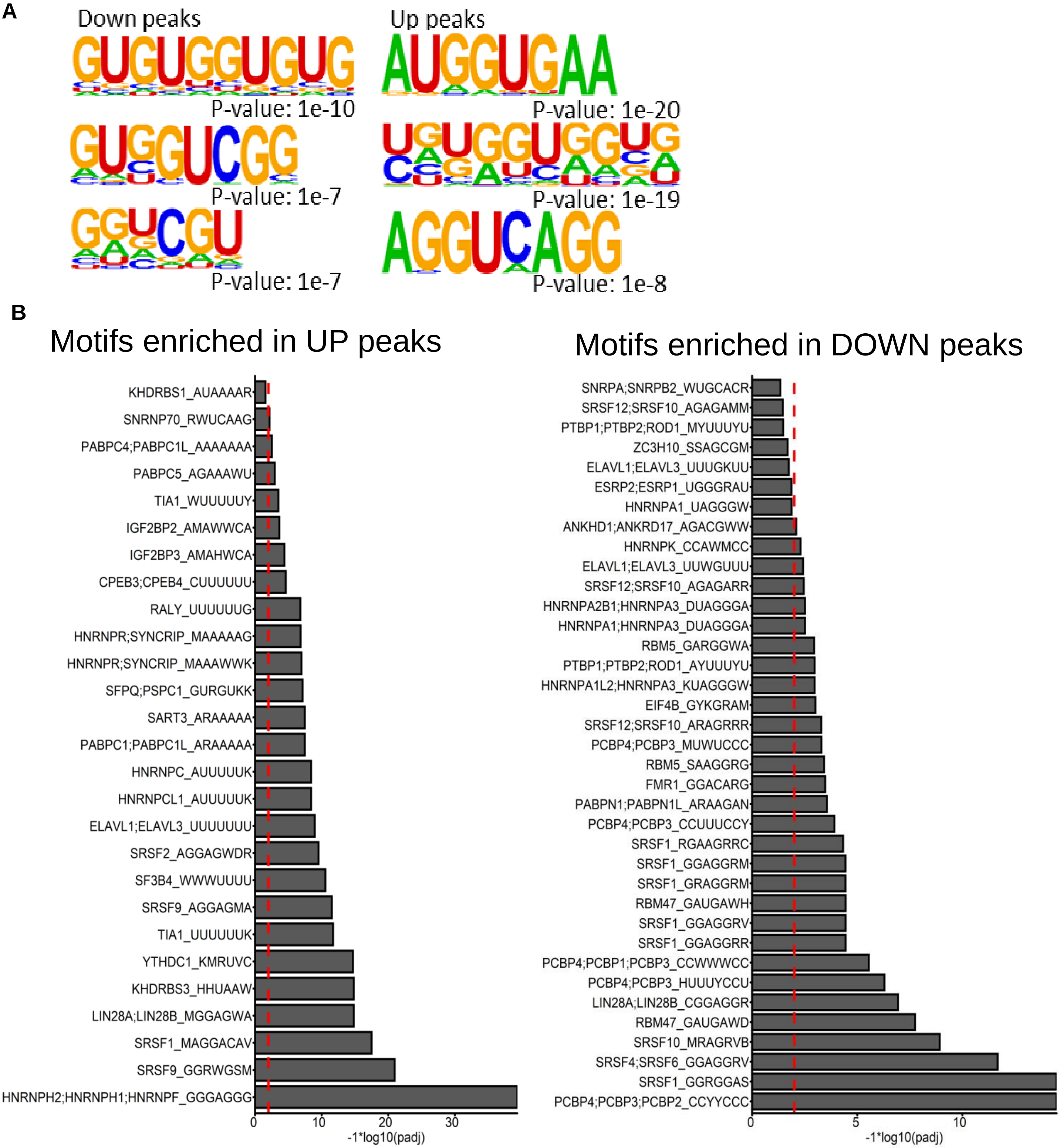


**Figure S7. Motif analysis of ePRINT peaks**

**A)** FUS-related motifs enriched in peaks that are reduced (UP peaks) or enhanced (DOWN peaks) in amplitude after FUS knockdown.

**B)** All RBP motifs identified as enriched in peaks that are enhanced (UP) or reduced (DOWN) in amplitude after FUS knockdown (adjusted p-value threshold of 0.05). Red dotted line indicates a p-value threshold of 0.01. Associated with Fig.1I.

Figure S8

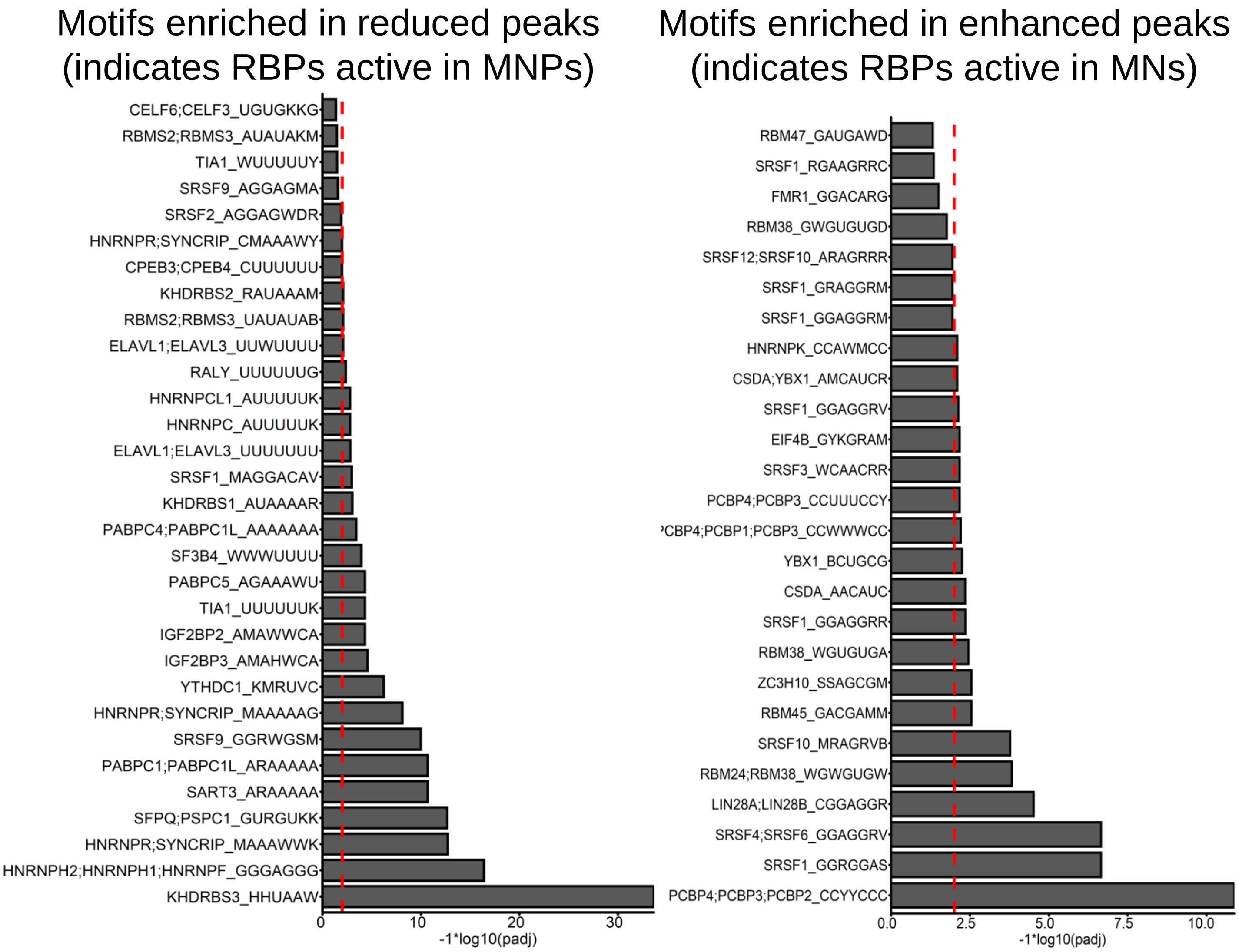


**Figure S8.** All RBP motifs identified as enriched in peaks that are enhanced or reduced in amplitude in MNs compared to MNPs (adjusted p-value threshold of 0.05). Red dotted line indicates a p-value threshold of 0.01. Associated with Fig.2D.

### Figure S9

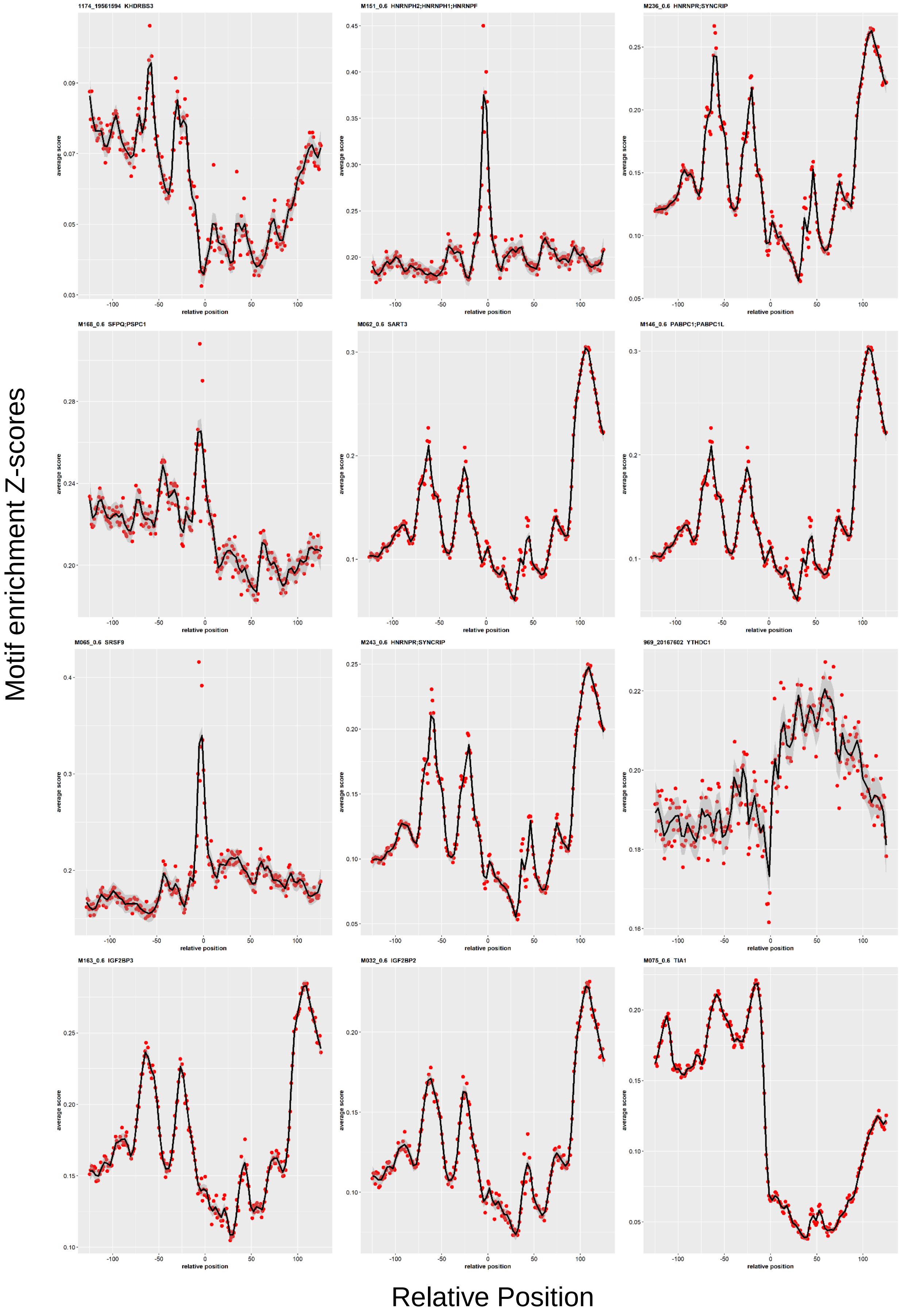


**Figure S9.** Motif location analysis to determine most probable binding location of RBPs with associated peaks reduced in MNs. A sharp peak at 0 indicated enrichment of binding location at peak start site.

### Figure S10

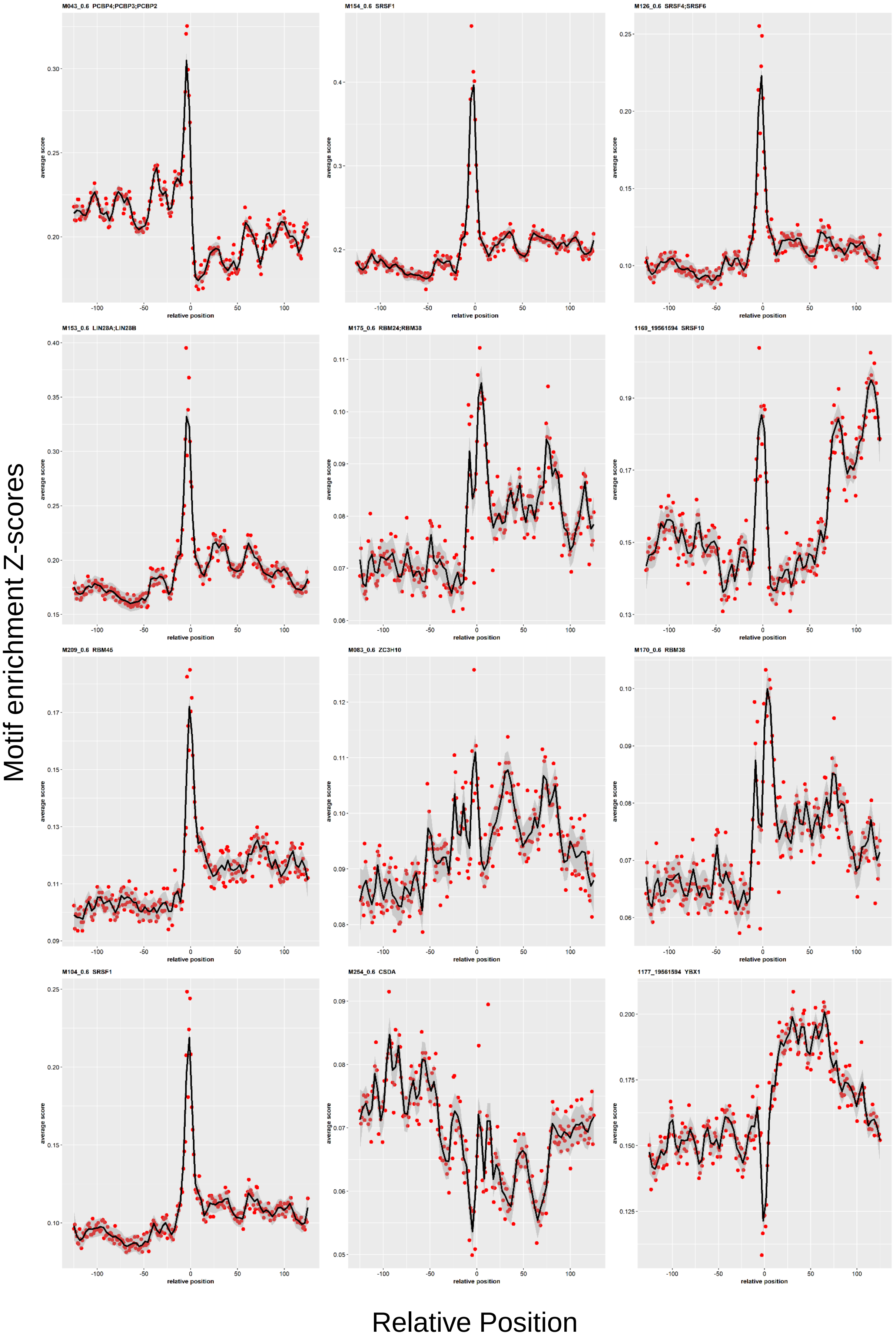


**Figure S10.** Motif location analysis to determine most probable binding location of RBPs with associated peaks enhanced in MNs. A sharp peak at 0 indicated enrichment of binding location at peak start site.

### Figure S11

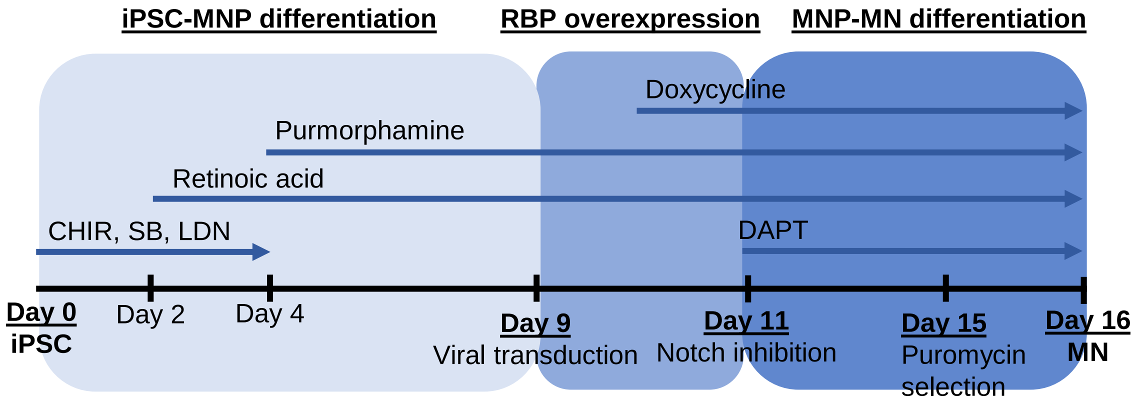


**Figure S11.** Experimental timeline for RBP overexpression during motor neuron differentiation. RBPs were expressed using dox-inducible viral vectors.

Figure S12

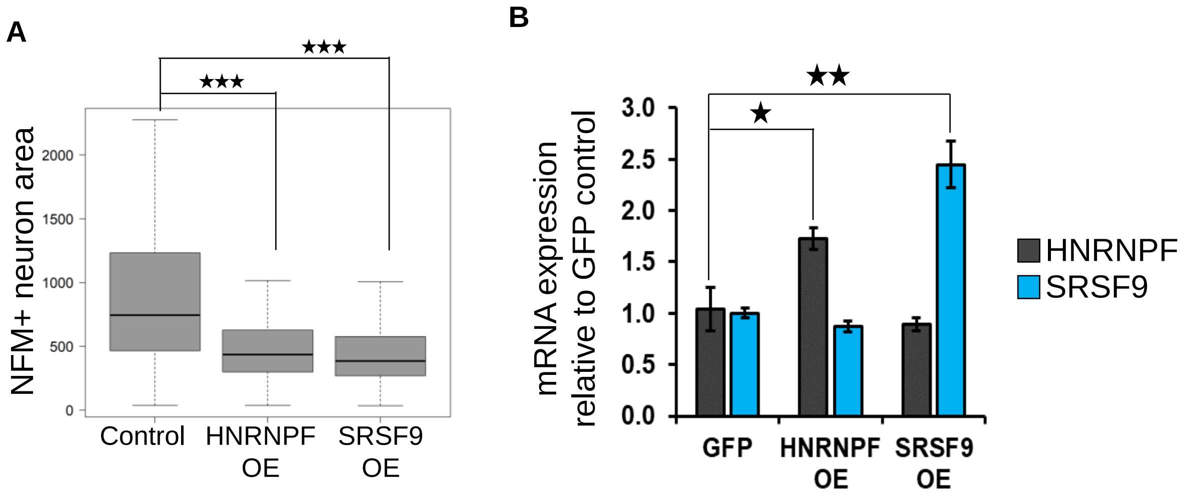


**Figure S12. Overexpression of HNRNPF and SRSF9 in motor neuron progenitors inhibits neurite outgrowth. Related to figure 2F**

**A)** Quantification of neuron size, measured by the area staining positive for NFM for each cell.

**B)** RT-qPCR validation of the overexpression constructs in HEK293 cells. Cells were transduced with dox-inducible and rtTA expressing lentiviruses and incubated for 48 hours in doxycycline-containing media after transduction. Transduction efficiency was ~50%. No antibiotic selection was performed before harvesting cells for RNA. N = 3, error bars indicate SEM.

* indicates pval <0.05. ** indicates pval <0.01. *** indicates pval <0.001 by Student's t-test.

### Figure S13

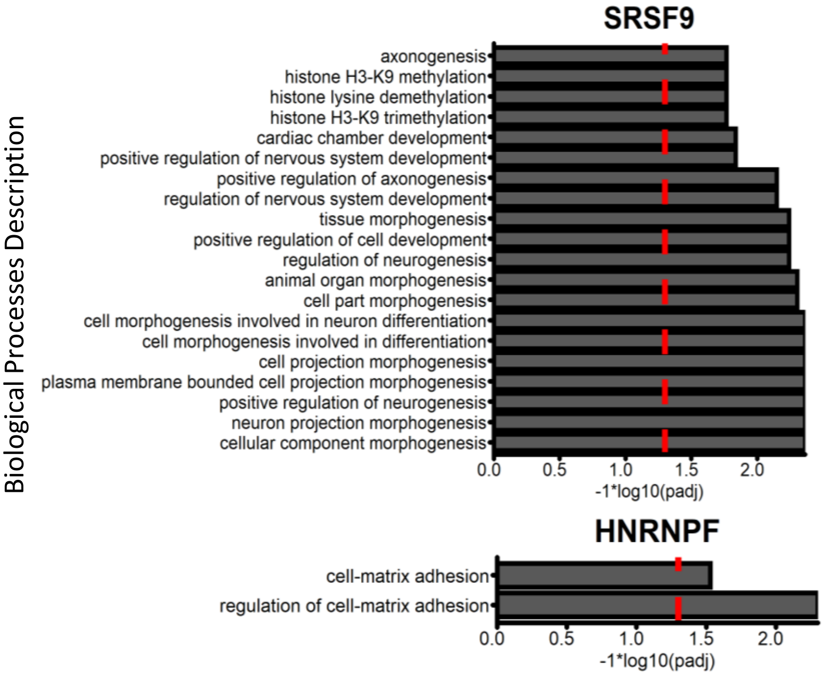


**Figure S13.** Gene ontology analysis of HNRNPF and SRSF9 target genes. Number of genes linked to a HNRNPF binding site = 623. Number of genes linked to an SRSF9 binding site = 857. Red line indicates an adjusted p-value of 0.05. Top 20 results shown for SRSF9 (total 34 enriched processes).

| symbol | log2FoldChange | padj |
|---|---|---|
| FUS | -1.178 | 4.43E-18 |
| TOMM20 | 0.731 | 1.53E-07 |
| MKNK2 | -0.967 | 3.16E-06 |
| KSR2 | -1.079 | 4.54E-06 |
| MIR2052HG | 0.838 | 5.74E-06 |
| HEY1 | 0.848 | 6.47E-06 |
| CA2 | -0.728 | 6.38E-05 |
| CA8 | -0.889 | 1.27E-04 |
| CIRBP | 0.777 | 1.30E-04 |
| ST8SIA5 | 2.004 | 1.72E-03 |
| PLS3 | 0.473 | 2.58E-03 |
| MCFD2 | 0.788 | 3.26E-03 |
| RPL34 | 0.675 | 5.37E-03 |
| DBN1 | -0.657 | 5.52E-03 |
| E2F2 | 0.771 | 7.38E-03 |
| HLCS | 0.540 | 8.64E-03 |
| CALU | 0.646 | 8.90E-03 |
| LINC00894 | 0.552 | 1.07E-02 |
| ALDH8A1 | 1.292 | 1.07E-02 |
| SLC7A5 | 0.584 | 1.07E-02 |
| TM7SF2 | -1.109 | 1.16E-02 |
| COL2A1 | -0.780 | 1.21E-02 |
| NSMF | 0.656 | 1.74E-02 |
| SLC6A16 | -1.039 | 1.74E-02 |
| RPL38 | 0.523 | 1.74E-02 |
| FRMD5 | -0.510 | 1.74E-02 |
| ISOC1 | 0.791 | 1.91E-02 |
| ARRDC4 | 1.236 | 1.91E-02 |
| GRB10 | 0.585 | 1.92E-02 |
| IRS4 | 0.545 | 1.92E-02 |
| SGPP2 | -1.120 | 1.97E-02 |
| NKAIN2 | -0.624 | 2.12E-02 |
| TMEM59L | -1.717 | 2.12E-02 |
| STMN1 | 0.513 | 2.12E-02 |
| SH3BP5-AS1 | 0.984 | 2.16E-02 |
| MIR3648-1 | 0.751 | 2.20E-02 |
| NEBL | -0.504 | 2.36E-02 |
| SRPRA | 0.637 | 2.36E-02 |
| NDRG4 | -0.841 | 2.39E-02 |
| EIF4G3 | 0.403 | 2.44E-02 |
| COLEC12 | -0.557 | 3.02E-02 |
| FRMD3 | 0.728 | 3.42E-02 |
| ID4 | 0.824 | 3.42E-02 |
| CAMK2B | -1.012 | 3.51E-02 |
| CSNK1E | 0.431 | 3.66E-02 |
| FARP2 | 0.537 | 3.66E-02 |
| GABPB2 | 0.625 | 3.85E-02 |
| DDX18 | 0.563 | 3.87E-02 |
| HAND1 | -0.972 | 4.12E-02 |
| GREB1 | -0.493 | 4.12E-02 |
| LRP2 | -0.490 | 4.49E-02 |
| SLC38A1 | 0.593 | 4.49E-02 |
| SCD5 | -0.543 | 4.68E-02 |

**Supplementary Table 1:** Differentially expressed genes (FUS knockdown in HEK293T cells)

| symbol | log2FoldChange | padj |
|---|---|---|
| ZFP36L1 | -5.526 | 6.22E-31 |
| RPS2 | -3.029 | 4.77E-20 |
| MBNL3 | -4.625 | 1.46E-16 |
| RAVER1 | -2.652 | 1.68E-15 |
| PTBP1 | -2.412 | 2.15E-14 |
| RPS19 | -1.940 | 2.97E-13 |
| RPLP0 | -2.204 | 1.10E-12 |
| RPS3 | -1.792 | 5.87E-12 |
| IGF2BP1 | -1.664 | 1.41E-11 |
| EEF1D | -2.099 | 3.65E-11 |
| RPL13A | -2.382 | 4.63E-11 |
| RBPMS | -2.323 | 7.49E-11 |
| MOV10 | -7.651 | 4.44E-10 |
| DNMT1 | -1.647 | 6.40E-10 |
| PARP4 | -3.031 | 6.98E-10 |
| SNRPB | -2.143 | 9.97E-10 |
| RPLP2 | -1.980 | 1.18E-09 |
| YBX3 | -3.286 | 1.76E-09 |
| RPL10A | -2.051 | 4.03E-09 |
| BRCA1 | -2.290 | 8.36E-09 |
| CELF2 | -1.911 | 1.09E-08 |
| RPL7A | -1.762 | 1.66E-08 |
| RPS6 | -1.428 | 1.86E-08 |
| RPL36 | -1.615 | 3.57E-08 |
| EXO1 | -7.071 | 4.73E-08 |
| RPL35 | -1.449 | 1.68E-07 |
| PARP1 | -1.466 | 2.32E-07 |
| CPSF1 | -1.485 | 2.65E-07 |
| RPL8 | -1.259 | 2.94E-07 |
| IPO4 | -1.695 | 3.13E-07 |
| RBM15 | -1.734 | 3.86E-07 |
| EIF3B | -1.454 | 4.23E-07 |
| ZFP36L2 | -3.950 | 5.62E-07 |
| RPS16 | -1.434 | 1.46E-06 |
| PRPF19 | -1.830 | 1.61E-06 |
| DHX38 | -1.955 | 1.84E-06 |
| FBL | -2.118 | 2.03E-06 |
| KPNB1 | -1.220 | 2.44E-06 |
| HNRNPUL1 | -1.426 | 3.27E-06 |
| GAPDH | -1.139 | 3.55E-06 |
| HNRNPM | -1.354 | 4.66E-06 |
| FUS | -1.237 | 7.88E-06 |
| KHSRP | -1.261 | 7.98E-06 |
| EIF4G1 | -1.148 | 1.06E-05 |
| SRSF2 | -1.755 | 1.57E-05 |
| RPL27 | -1.481 | 1.89E-05 |
| SETD1A | -1.389 | 2.36E-05 |
| EFTUD2 | -1.501 | 2.97E-05 |
| RPLP1 | -1.575 | 3.02E-05 |
| RBPMS2 | -3.446 | 4.46E-05 |
| SRRT | -1.388 | 4.61E-05 |
| SRRM2 | -1.078 | 5.94E-05 |
| RNASEH2A | -2.663 | 8.04E-05 |
| ALYREF | -1.568 | 8.88E-05 |
| HNRNPAB | -1.345 | 9.34E-05 |
| RPS8 | -1.472 | 9.35E-05 |
| RPL32 | -0.876 | 9.78E-05 |
| RPL18 | -1.658 | 1.03E-04 |
| RPS5 | -1.329 | 1.14E-04 |
| LSM4 | -1.282 | 1.22E-04 |
| U2AF1 | -1.367 | 1.34E-04 |
| SNRNP200 | -1.075 | 1.65E-04 |
| RPL28 | -1.235 | 1.71E-04 |
| SART3 | -1.251 | 2.06E-04 |
| HNRNPF | -1.573 | 2.32E-04 |
| POLDIP3 | -1.267 | 2.38E-04 |
| SARS2 | -2.201 | 2.57E-04 |
| SART1 | -1.265 | 2.80E-04 |
| RPL37A | -0.964 | 3.47E-04 |
| BICC1 | -1.975 | 4.91E-04 |
| PRPF8 | -0.940 | 5.15E-04 |
| INTS1 | -1.113 | 5.30E-04 |
| RPL19 | -1.120 | 5.32E-04 |
| RPL23A | -2.212 | 5.33E-04 |
| DDX54 | -1.318 | 6.54E-04 |
| RBM15B | -1.224 | 7.24E-04 |
| DDX23 | -1.277 | 7.24E-04 |
| HNRNPU | -0.859 | 7.70E-04 |
| HDLBP | -0.914 | 9.51E-04 |
| RPL13 | -1.073 | 9.77E-04 |
| SON | -0.898 | 1.03E-03 |
| EIF3G | -1.426 | 1.04E-03 |
| PCBP2 | -1.066 | 1.32E-03 |
| PABPC1 | -0.932 | 1.38E-03 |
| CNOT3 | -1.244 | 1.45E-03 |
| DGCR8 | -1.097 | 1.46E-03 |
| DHX37 | -1.705 | 1.65E-03 |
| RPS11 | -0.864 | 1.73E-03 |
| SRRM1 | -1.061 | 1.76E-03 |
| RPS4X | -0.813 | 1.87E-03 |
| RPL27A | -1.380 | 1.92E-03 |
| RBM10 | -0.994 | 1.94E-03 |
| RPS15 | -1.754 | 2.01E-03 |
| DDX46 | -1.079 | 2.22E-03 |
| GEMIN8 | -1.889 | 2.35E-03 |
| SMG5 | -1.275 | 2.36E-03 |
| PRPF31 | -1.475 | 2.41E-03 |
| RPS14 | -1.151 | 2.82E-03 |
| SRSF1 | -0.979 | 2.92E-03 |
| SF1 | -0.892 | 3.01E-03 |
| U2AF2 | -0.974 | 3.04E-03 |
| EEF2 | -0.796 | 3.70E-03 |
| RRS1 | -2.573 | 3.75E-03 |
| HNRNPH3 | -1.006 | 3.90E-03 |
| RRP1 | -1.412 | 4.17E-03 |
| NUTF2 | -1.489 | 4.26E-03 |
| TBRG4 | -1.763 | 4.37E-03 |
| RPS9 | -1.129 | 4.74E-03 |
| RPL23 | -0.931 | 4.90E-03 |
| PUS1 | -1.641 | 4.95E-03 |
| EIF4A3 | -0.945 | 5.25E-03 |
| UPF1 | -1.014 | 5.53E-03 |
| G3BP1 | -1.061 | 6.16E-03 |
| EEF1A1 | -1.243 | 6.59E-03 |
| EEF1G | -1.124 | 6.62E-03 |
| GTPBP1 | -1.096 | 7.40E-03 |
| SAFB | -0.863 | 7.57E-03 |
| EIF3A | -0.886 | 7.74E-03 |
| SRSF9 | -1.096 | 8.39E-03 |
| MRPL4 | -2.065 | 8.55E-03 |
| RBMX | -0.909 | 9.12E-03 |
| SF3A1 | -1.085 | 1.01E-02 |
| RPL11 | -0.867 | 1.05E-02 |
| CPSF3 | -1.192 | 1.11E-02 |
| RPS15A | -1.255 | 1.34E-02 |
| ZCCHC3 | -1.251 | 1.34E-02 |
| UTP20 | -1.269 | 1.35E-02 |
| EIF5A | -1.021 | 1.43E-02 |
| FBXO17 | -2.409 | 1.46E-02 |
| PRPF38A | -1.245 | 1.49E-02 |
| GEMIN4 | -1.493 | 1.63E-02 |
| RBM19 | -0.925 | 1.70E-02 |
| LARP7 | -1.405 | 1.77E-02 |
| SRP68 | -1.158 | 1.80E-02 |
| DDX39A | -1.456 | 1.87E-02 |
| CNOT1 | -0.816 | 2.11E-02 |
| HNRNPL | -0.800 | 2.12E-02 |
| SF3B2 | -0.720 | 2.16E-02 |
| MRPL11 | -2.798 | 2.22E-02 |
| RRBP1 | -0.864 | 2.52E-02 |
| RRP7A | -1.312 | 2.60E-02 |
| CPSF7 | -0.921 | 2.66E-02 |
| MRPS34 | -1.151 | 2.74E-02 |
| MRPS18A | -1.830 | 2.80E-02 |
| EXOSC6 | -1.648 | 2.89E-02 |
| RPL30 | -0.952 | 3.01E-02 |
| RPS20 | -0.881 | 3.15E-02 |
| LSM14B | -0.938 | 3.20E-02 |
| EIF4H | -0.960 | 3.35E-02 |
| PUF60 | -1.173 | 3.49E-02 |
| HNRNPC | -0.647 | 3.51E-02 |
| SNRNP25 | -1.738 | 3.56E-02 |
| NCL | -0.604 | 3.60E-02 |
| RPL39L | -3.567 | 3.89E-02 |
| UTP14A | -1.374 | 4.00E-02 |
| SLBP | -1.290 | 4.01E-02 |
| SF3B3 | -0.724 | 4.04E-02 |
| PABPC4L | -7.176 | 4.22E-02 |
| SPATS2L | -1.239 | 4.28E-02 |
| ATXN2L | -0.776 | 4.49E-02 |
| SF3A2 | -0.917 | 4.85E-02 |
| RBMS1 | -0.885 | 4.88E-02 |
| SF3B4 | -1.344 | 4.93E-02 |

**Supplementary table 2:** Differentially expressed RBP genes (MNP to MN), ordered by p-value. HNRNPF and SRSF9 are highlighted in red.
